# Distinguishing biological from technical sources of variation by leveraging multiple methylation datasets

**DOI:** 10.1101/521146

**Authors:** Mike Thompson, Zeyuan Johnson Chen, Elior Rahmani, Eran Halperin

**Affiliations:** Department of Computer Science, University of California Los Angeles, Los Angeles, CA, USA; Department of Human Genetics, University of California Los Angeles, Los Angeles, CA, USA; Department of Anesthesiology and Perioperative Medicine, University of California Los Angeles, Los Angeles, CA, USA; Department of Biomathematics, University of California Los Angeles, Los Angeles, CA, USA

## Abstract

DNA methylation remains one of the most widely studied epigenetic markers. One of the major challenges in population studies of methylation is the presence of global methylation effects that may mask local signals. Such global effects may be due to either technical effects (e.g., batch effects) or biological effects (e.g., cell-type composition, genetics). Many methods have been developed for the detection of such global effects, typically in the context of epigenome-wide association studies. However, current unsupervised methods do not distinguish between biological and technical effects, resulting in a loss of highly relevant information. Though supervised methods can be used to estimate known biological effects, it remains difficult to identify and estimate unknown biological effects that globally affect the methylome. Here, we propose *CONFINED,* a reference-free method based on sparse canonical correlation analysis that captures replicable sources of variation—such as age, sex, and cell-type composition—across multiple methylation datasets and distinguishes them from dataset-specific sources of variability (e.g., technical effects). Consequently, we demonstrate through simulated and real data that by leveraging multiple datasets simultaneously, our approach captures several replicable sources of biological variation better than previous reference-free methods and is considerably more robust to technical noise than previous reference-free methods. *CONFINED* is available as an R package as detailed at https://github.com/cozygene/CŪNFINED.

## 1 Introduction

While technological advances have provided a surplus of methylation datasets, analyses of these datasets are often complicated by innumerable possible sources of variability [1,2]. In particular, epigenome-wide association studies (EWAS) and studies that aim to implicate observed methylation signal to phenotypic variance are particularly at risk for false associations due to unknown drivers of the observed signal that globally affect the epigenome [3–5]. For example, age is correlated with a large number of methylation sites and phenotypes [6–8], and thus if not corrected for, association between a specific methylation site and a phenotype may be primarily driven by a confounder such as age. In order to mitigate spurious associations in such association studies, it is crucial to elucidate and account for the sources of variation that globally affect the methylation patterns in the genome.

Sources of global methylation effects can be either technical or biological, and may also be measured or unmeasured. In the case of technical sources, most typical are batch effects, or variation resulting from different technicians or conditions during the data-preparing steps [9]. These sources should undoubtedly be identified and accounted for in analyses, for example by balancing cases, controls, and samples from different datasets, including measured potential confounders as covariates, regressing out the sources of confounding signals if they are measured, or otherwise estimating these potential sources of technical effects and accounting for their estimates [10].

The case of biological sources is more complex; biological sources of variation such as age, sex, cell-type composition, genetics, ethnicity, co-morbidities, or responses to environmental factors like medication intake or smoking status indeed affect the global methylation patterns in the genome, and they are also often correlated to the phenotype of interest [6,11–15]. However, due to logistical limitations, often only a few of these sources of biological variation are measured in a given study; moreover, it is often the case that, some of the sources of variation that are correlated with the phenotype are unknown and hence unmeasured.

Unlike technical effects, there is much debate over the best practice of using these biological sources of variation in a model (e.g., [3,13,16,17]) since one can argue that identifying these sources is an important ingredient in understanding the disease mechanism. Moreover, identifying these biological sources of variation may be useful in prediction algorithms related to the studied phenotype. In other words, it is context-specific whether one should include biological sources of variation in their model—considering the additional sources as confounders—or simply derive a model considering only the observed signal and accounting for the technical effects [18].

To capture signal corresponding to specific biological sources of variation, reference-based methods have been proposed. In the case of methylation, one commonly researched source of biological variability is cell-type composition. Houseman et al. developed an approach to estimate the true cell-type proportions in methylation datasets using “methylation signatures” (estimates of cell-type-specific methylation levels across a population) [19]. Reference-based methods and methods that leverage prior statistics, however, are limited to known sources of variability for which such reference data exists. In many cases, either the sources of variability are unknown, or there is no reference data that can be utilized for these methods (e.g., factors such as diet and exposure to air pollution [20–22], and tissues such as solid tumors or adipose [23]). In such cases, reference-based methods cannot be used.

In an attempt to overcome the above limitations, many reference-free methods [23–29], have been proposed. Though these methods can correct for cell-type composition in EWAS [27,30] and may also capture other sources of variability, they are limited by the fact that it is impossible to know whether their components reflect biological or technical signal (Figure 1). While technical signal is not of interest and should be accounted for in the analysis, the biological signal can provide useful insights about, underlying biological phenomena., for instance by being used to model the interaction with the methylation signal.

**Figure 1.**
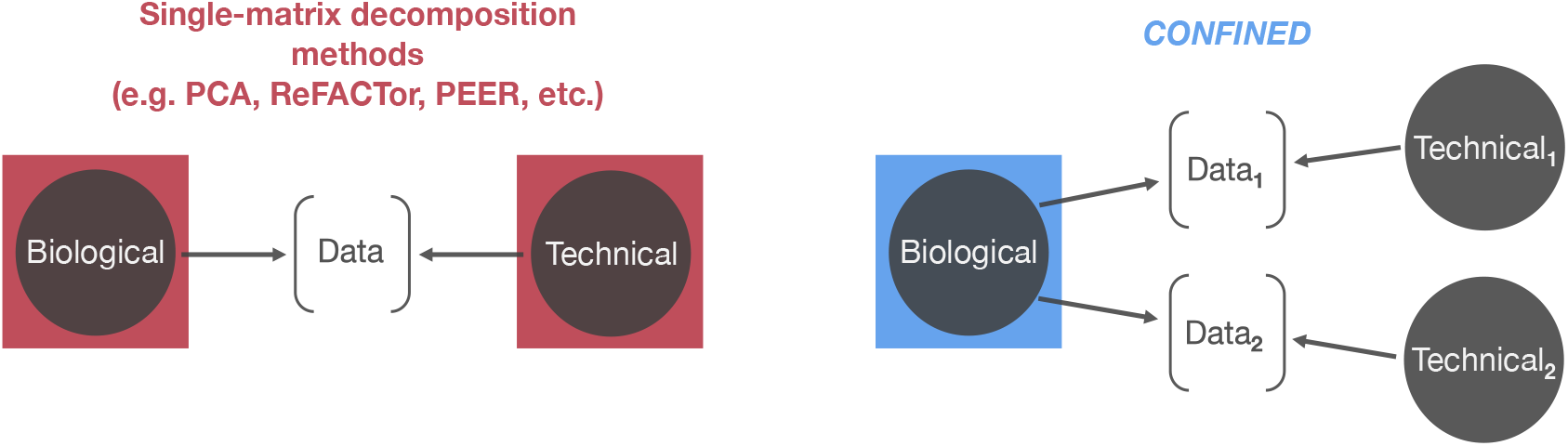
*CONFINED* compared to previous factorization approaches. Previous reference-free methods based on single-matrix decompositions (e.g. principal component, analysis, non-negative matrix factorization) capture the dominant, sources of variability which may be composed of both biological and technical effects (left). Here, we propose a. method to capture solely biological variability (right).

In this paper, we propose a reference-free method that disentangles the technical sources of variation from the biological sources of variation. Our method is based on the observation that the same biological sources of variation typically affect different studies that are performed under the same conditions (e.g., on the same tissue type), while technical variability is study-specific. Thus, unlike previous unsupervised methods that utilize single-matrix decomposition techniques to account for covariates in methylation data, we propose the use of canonical correlation analysis (CCA), which captures shared signal across multiple datasets. In brief, CCA finds shared structure between two datasets by finding maximally-correlated linear transformations of the datasets and is used across many fields including cognitive science [31], psychology [32], and imaging [33]. CCA has been used in the context of genomics to capture genome-wide similarities between different genomic measurements (e.g., gene expression and genetics [34,35], gene expression and copy number alterations [36,37]) for the same set of individuals. As opposed to this traditional use of CCA, our method, named *CONFINED* (CCA ON Features for INter-dataset Effect Detection), searches for genome-wide similarities between one methylation profile across two sets of individuals. By instead searching across a single genomic profile, we capture shared structure inherent to the underlying biology of the datasets.

We evaluated the performance of *CONFINED* through both simulated and real data. Our evaluations demonstrate that *CONFINED* captures signal from only biologically replicable sources of variability. We show, as examples, improvement over previous methods by comparing their performance in capturing methylation signal due to cell-type composition, age, and sex in several whole-blood datasets. We also demonstrate that by inducing sparsity, *CONFINED* prioritizes features that recapitulate biological functionality inherent to both datasets. For example, when pairing two whole-blood datasets together, the sites best ranked by *CONFINED* were significantly enriched for immune cell function.

## 2 Results

### A brief summary of *CONFINED*

We developed *CONFINED* to capture biological sources of variability in methylation datasets. As input, *CONFINED* takes two matrices with the same number of rows (methylation sites) but not necessarily the same number of columns (individuals), k the number of components to produce, and *t* the number of CpG sites to use, or in other words, a sparsity parameter. As output, *CONFINED* produces k components that can be used to model biological sources of variability for each input dataset.

Notably, *CONFINED* is based on CCA which considers two datasets simultaneously. Intuitively, CCA performs a decomposition of two matrices simultaneously, and hence finds linear combinations of features that define biological variation present in both datasets. Conversely, previous methods that decompose one matrix at a time essentially look for linear or non-linear (kernel-based) combinations of features that preserve dominant structure in a single dataset, and this structure may be a combination of both biological and technical signal. Thus, leveraging the shared structure of two datasets through CCA is crucial. Nonetheless, there are two substantial differences between *CONFINED* and traditional uses of CCA in genomic studies. First, *CONFINED* looks for shared structure of one methylation profile across two sets of individuals rather than looking for shared structure in one set of individuals across two sets of genomic measurements. Second, *CONFINED* performs a feature selection procedure that is critical to detect the shared sources of variability across the different datasets.

### *CONFINED* distinguishes between technical and biological signal: Real data analysis with simulated batch effects

In the context of capturing biological signal, one of the main limitations of single-matrix decomposition methods (e.g., PCA, ReFACTor [24], PEER [38], non-negative matrix factorization (NNMF) [39]), is that each of their components may consist of a mixture of signal reflective of technical noise specific to a dataset, such as batch effects, and the biological signal. For instance, PCA and methods based on PCA, such as ReFACTor [24] and penalized matrix decomposition (PMA) [36], consider directions in the data that explain the most variability, but this variability is not limited to strictly global biological or replicable effects in the individual datasets. This issue may also be present in PEER [38], which includes a probabilistic version of factor analysis, as the latent factors driving the data may also include some effect from technical variability. Similarly, in NNMF [39] a data matrix is decomposed as a linear combination of different components, and some of the signal of the data matrix may be deconstructed by a component that captures technical variation. Intuitively, *CONFINED* should be robust to dataset-specific technical effects as it only looks for shared structure across datasets.

To illustrate that *CONFINED* captures only replicable biological signal, we simulated batch effects for two whole-blood methylation datasets from Hannum et al. [41] and Liu et al. [40] and compared our method to several earlier methods based on single-matrix decomposition. In this setting, we generated dataset-specific noise with low-rank structure and added it to each of the datasets prior to running any feature selection or method. Naturally, simulated batch effects induce technical variation in the datasets, and thus may interfere with methods’ abilities to capture biological variation. We used the datasets with added noise to capture cell-proportion estimates of the original datasets as reported by the method proposed by Houseman et al. [19]. Houseman et al. proposed a reference-based method for estimating proportions of immune cells in whole-blood methylation data by leveraging differentially methylated regions of DNA to form methylation signatures for individual cell-types. They then use these signatures to obtain cell proportion estimates for several immune cells (CD4 *t* cells, CD8 *t* cells, B cells, natural killer cells, monocytes and granulocytes). In our experiments, we fit a linear model of each Houseman-estimated cell-type proportions using several components from each of the methods (Figure 2).

**Figure 2.**
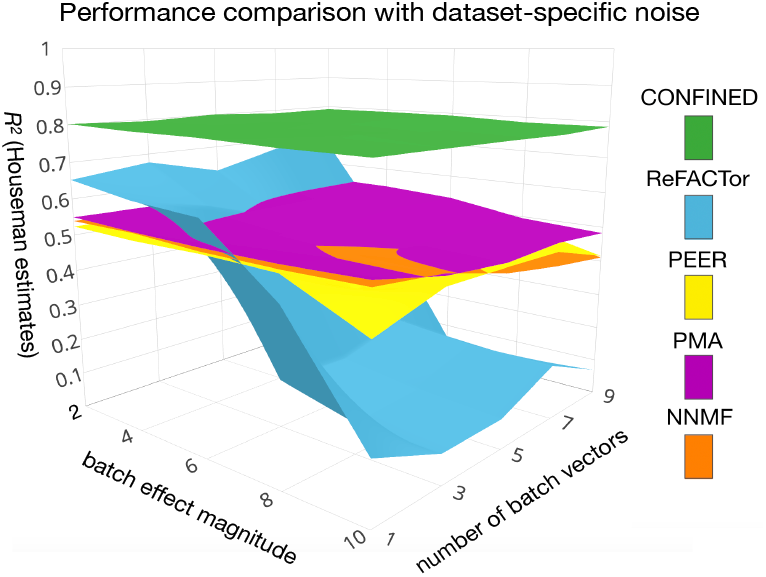
Capturing cell-composition in the presence of simulated technical noise. We added simulated batch effects to the whole-blood datasets of Liu et al. [40] and Hannum et al. [41] and compared the ability of *CONFINED*, ReFACTor [24], PEER [38], PMA [36], and NNMF to capture cell-type composition in whole-blood. Here, we show the results of the Hannum et al. dataset, however the results of each method were quantitatively similar across both datasets (Supplementary Figure S9).

We evaluated the performance of each method while varying the strength of simulated, dataset-specific technical effects (Methods, Supplementary Methods; Section S5). The components of *CONFINED* best captured the biological signal and were the only components that were robust to technical variation across all levels of noise (Figure 2). In addition to the biological signal, the components of the previous methods captured signal pertaining to the simulated batch effects (Supplementary Methods; Section S5).

### *CONFINED* finds biological sources of variability with high accuracy: Analysis across multiple real datasets

We evaluated *CONFINED* using the whole-blood methylation datasets from Hannum et al. [41] and Liu et al. [40] as well as a dataset from Lunnon et al. [42] containing brain tissue samples. Along with their methylation data were measured sources of biological variation including patients’ disease status, age, sex and location from which the brain sample was taken. In addition to evaluating *CONFINED*’s ability to capture the measured biological factors, we also evaluated its performance on an unmeasured source of variation, cell-type composition. While we focused on using two datasets corresponding to the same tissue type in several analyses, we note that the studied phenotypes in the datasets were different (e.g., Hannum et al. studied aging whereas Liu et al. studied Rheumatoid arthritis). In these experiments, we considered only known shared biological sources of variation and therefore excluded from our analyses sources of variation that may only appear in one dataset, e.g. patient status or brain location. As we show below, using *CONFINED* we were able to produce components that correlated with both the measured and unmeasured sources of biological signal across all datasets.

First, we evaluated *CONFINED* against other reference-free methods when capturing unmeasured biological sources of variability in two whole-blood datasets. In one experiment, we used *CONFINED* to capture cell-type composition, which was unmeasured in both studies. We treated cell-type proportion estimates from the reference-based algorithm of Houseman et al. [23] as the ground-truth. *CONFINED* outperformed all of the previous methods we tested, with pronounced differences in its estimation of the composition of monocytes and natural killer cells (Figure 3, Supplementary Figure S4). Additionally, we considered the situation in which two datasets are concatenated and supplied to a single-matrix-decomposition method as a single dataset, as well as the situation in which a single-matrix decomposition method leverages the features selected by *CONFINED*. In both procedures, however, the components of the single-matrix method were less correlated to cell-type composition than the components of *CONFINED* (Supplementary Methods; Sections S4, S3).

**Figure 3.**
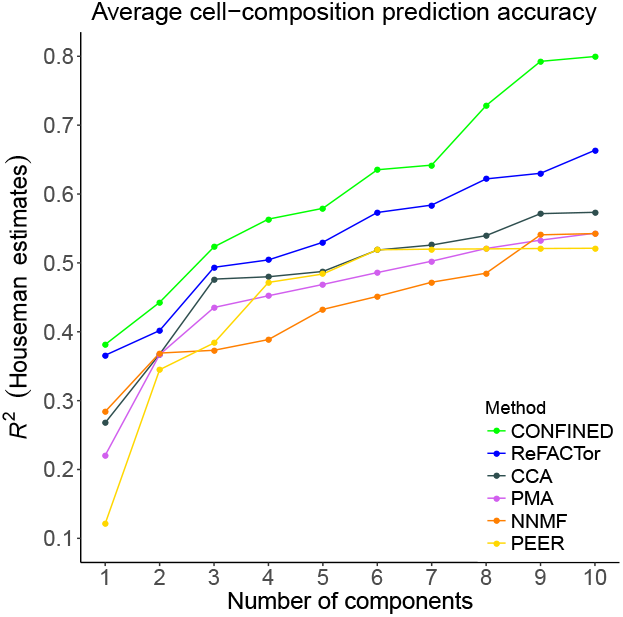
A comparison of *CONFINED* and previous reference-free methods in capturing leukocyte composition. We used each methods’ components to capture cell-type proportions as estimated by the reference-based method of Houseman et al. [19] across CD4 *t* cells, CD8 *t* cells, monocytes, B cells, natural killer cells, and granulocytes in whole-blood data from an aging study (Hannum et al. [41]) as well as in whole-blood from a study of Rheumatoid arthritis (Liu et al. [40] Supplementary Figure S5).

We also used *CONFINED’s* components to capture measured sources of biological variation across tissue-types (Figure 4). In these experiments, we paired a whole-blood dataset [40] first with another whole-blood dataset [41], and second, with a dataset composed from brain tissue [42]. Notably, the accuracy of *CONFINED* to capture each source of signal varied depending on the pairing of the tissue-type (i.e blood-blood vs. blood-brain) and the sparsity parameter used.

In the whole-blood dataset, *CONFINED*’s components captured age and sex with accuracy 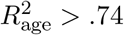 and 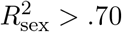 respectively (Figure 4). In the case of other methods, PMA [36] had the highest performance among previous methods, but was greatly outperformed by *CONFINED* (Supplementary Methods; Section S8). Notably, using relatively less sparsity to capture age and sex achieved greater accuracy, however this trend was not necessarily observed when using lower sparsity for capturing cell-type composition.

**Figure 4.**
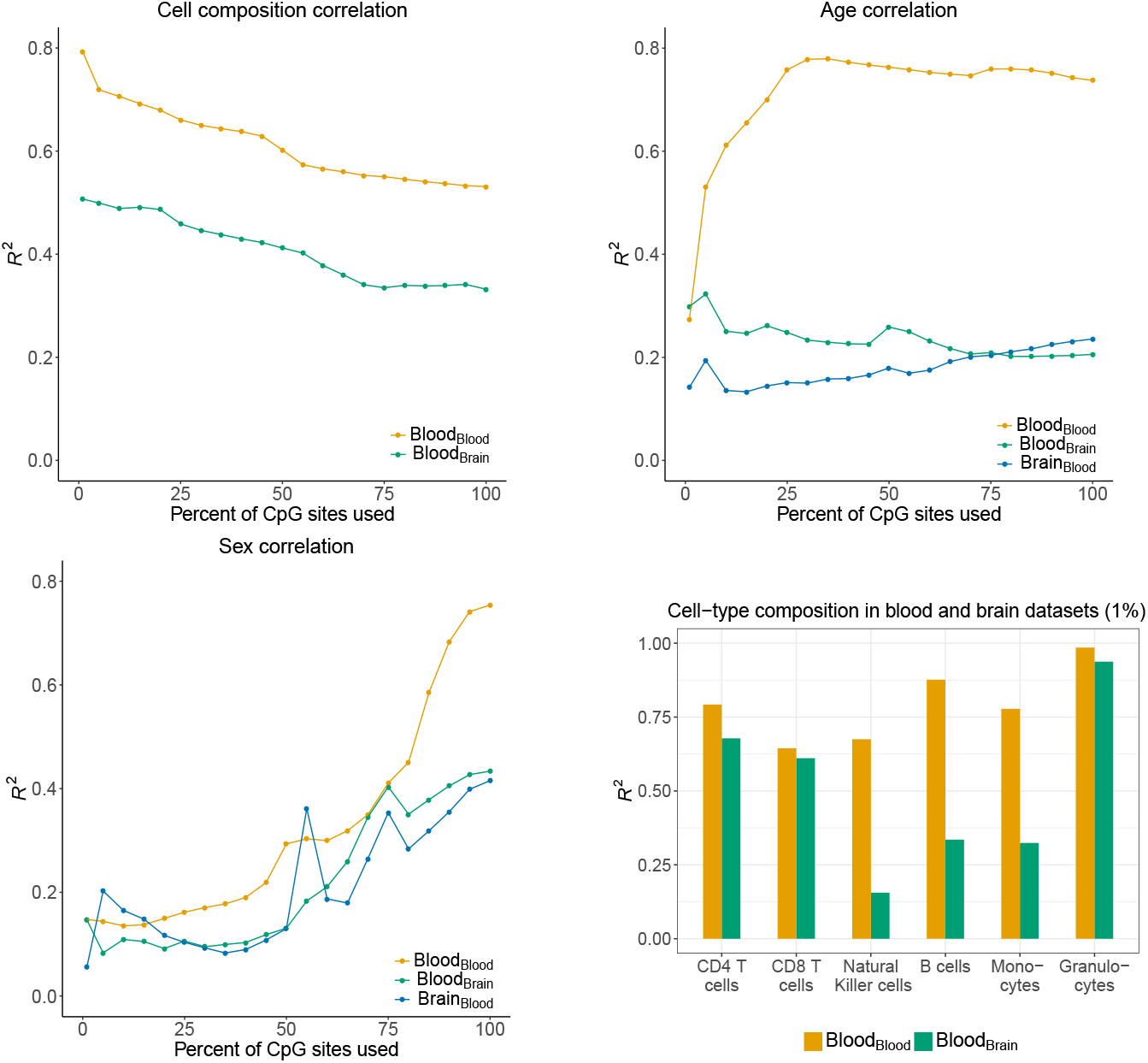
Biological drivers of variability captured by *CONFINED* across a range of sparsity. We paired a whole-blood dataset [40] with another whole blood dataset [41] (not shown) and with a brain dataset [42] to capture sources of variability in each dataset. The subscript indicates with which tissue-type the dataset was paired. We fit a linear model for each source of variability was using 10 *CONFINED* components to obtain an R^2^ value. We varied the percentage of CpG sites used from 1% (nearly entirely sparse) to 100% (no sparsity).

When pairing the blood dataset with the brain dataset, *CONFINED*’s components were correlated with some of the whole-blood dataset’s measured biological factors with slightly less strength than when pairing it with a dataset of the same tissue type 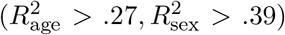 (Figure 4), possibly suggesting a different architecture for genome-wide variation across the different tissue types. Nonetheless, the cell-type composition accuracy for the blood dataset when paired with the brain dataset was still relatively high (average 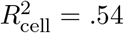). This is likely due to the fact that several types of immune cells are known to populate or have immune-related functions in the brain (e.g. resident *t* cells [43,44], glia [45] and neutrophils (granulocytes) [46]). Therefore, the immune function of cells in the brain and immune cells in the blood may follow similar pathways that could be reflected in the epigenome. The biological sources of variability in the brain dataset were captured with overall less accuracy than the whole-blood biological sources of variability 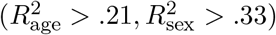.

### Gene ontology of *CONFINED*’s methylation sites

We evaluated the biological significance of the features selected by *CONFINED* using the R package `missMethyl` [47]. For a given set of methylation sites, `missMethyl` tests for enrichment in gene ontology (GO) pathways by first mapping the sites to genes (weighing the genes based on the number of sites that map to them), then performing a test built off of Wallenius’ noncentral hypergeometric distribution. In order to avoid potential biases resulting from the parametric assumptions in the model of `missMethyl`, we performed permutation testing using its reported p-values. Our test yielded significant enrichment for various ontologies across each pair of datasets (Table 1).

**Table 1.** Gene Ontology Enrichment of sites ranked by *CONFINED.* We tested enrichment of the highest-ranked sites by *CONFINED* in a blood-blood pair of datasets. Here, we set the sparsity parameter based on a rule learned through cross-validation (Supplementary Methods; Section S6), however we observed similar results across a range of sparsity parameters (Supplementary Methods; Section S7).

| Ontology term | p-value (permutation) | p-value (missMethyl) |
| --- | --- | --- |
| Immune system process | .001 | 6.9e−18 |
| Immune response | .001 | 1.0e−15 |
| Regulation of immune response | .026 | 3.0e−11 |

When we paired two whole-blood datasets, the highest ranked features by *CONFINED* were enriched for pathways generally involved with the immune response, leukocyte activation, and defense response. Notably, most of the significantly enriched pathways were related to the immune system or signaling (Table 1). These results underscore the importance of *CONFINED*’s sparsity and provide support for *CONFINED*’s ability to capture biologically meaningful signal.

Pairing the blood and brain datasets, we observed somewhat similar results, but with less significance. The most enriched pathways in the blood-brain pair included several immune system or hematopoietic processes, but the less enriched pathways were primarily different than when pairing the two blood datasets. The pathways in the blood-brain pair were generally not significantly enriched using permutation testing, unless we used a relatively lower level of sparsity.

## 3 Methods

### A brief explanation of canonical correlation analysis

We first explain the general idea of canonical correlation analysis (CCA) [48]. In the simplest terms, CCA maximizes the correlation of two matrices via linear transformations. CCA takes as input two matrices *X*_1_ of dimension *n* × *m*_1_ and *X*_2_ of dimension *n* × *m*_2_ where *n* > *m*_1_ and *m*_2_. In other words, both matrices have the same number of rows, but not necessarily the same number of columns. CCA then attempts to find *m*_1_ and *m*_2_-length vectors *a*_1_ and *a*_2_, such that the correlation of *X*_1_a_1_ and *X*_2_*a*_2_ is maximized:

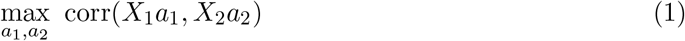

The solutions *a*_1_ and *a*_2_ are obtained from the first eigenvectors of subsequently generated matrices, as detailed by Hardoon et al. [49]. We define the products *X*_1_*a*_1_ and *X*_2_*a*_2_ as the first canonical variables and let *u*_1_ = *X*_1_*a*_1_ and *u*_2_ = *X*_2_*a*_2_. We can find additional pairs of canonical variables with subsequently less correlation under the constraint that each pair of canonical variables is orthogonal to each other. Notably, there are min{*m*_1_,*m*_2_} eigenvectors, thus there are min{*m*_1_,*m*_2_} canonical variables. We define the collection of canonical variables for each dataset as follows:

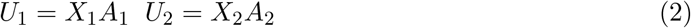

For completeness, we derive and include a tutorial for canonical correlation analysis in Supplementary Methods and Information section S1.

### A formal description of *CONFINED*

CCA has been used in genomics in many instances [50–52]. In these cases the rows correspond to individuals, while the columns correspond to features of genomic measurements. For example, each feature could be the expression of a specific gene in one matrix, and in the other matrix it could be the genotype allele, i.e., in this case *X*_1_ corresponds to a gene expression matrix, and *X*_2_ corresponds to a genotype matrix, but both measurements have been taken on the same set of individuals. In *CONFINED*, we transpose the problem. Rather than searching for shared directions between two sets of genomic measurements, we instead search for shared directions of the same type of genomic measurement (in our case, methylation), but across two sets of individuals. Moreover, since we find that in practice many sources of variability in methylation only act on a fraction of the methylation sites in the genome [14,24], *CONFINED* uses sparsity by limiting the analysis to a fraction of the methylation sites in the genome. We note that our method shares similarities with a recent application of CCA to single-cell expression datasets [53]. However, unlike this method, we search for shared structure across two sets of individuals rather than two sets of cells, and we assume the number of genomic features is larger than the number of individuals (or cells).

Formally, *CONFINED* takes as input two matrices, *X*_1_ with dimension *m* × *n*_1_ and *X*_2_ with dimension *m* × *n*_2_, of *m* measured methylation sites for *n*_1_ and *n*_2_ individuals respectively. In addition, it takes as input a sparsity parameter *t*, a dimensionality parameter *l*, and an output parameter specifying the number of components to generate *k*. To generate its components, *CONFINED* first selects the *t* most informative features then runs CCA on these *t* features:

1. Obtain *U*_1_ and *U*_2_ both of size *m* × min{*n*_1_,*n*_2_} following Equations (1) and (2).
2. Construct *Ũ*_1_ and *Ũ*_2_ both of dimension *m* × *l* from the first *l* columns of *U*_1_ and *U*_2_ respectively.
3. Generate a low-rank approximation of each dataset:

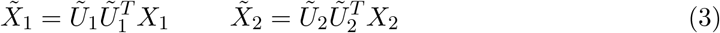
4. For each site *j* in dataset *i* compute a score based on its correlation between itself and its low-rank approximation:

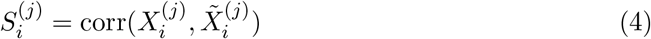
5. Rank the sites with the highest inter-dataset score:

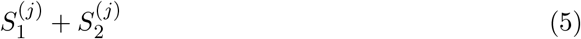
6. Perform CCA using the sites with the top *t* scores, returning *CONFINED* components 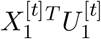 of size *n*_1_ × *k* for *X*_1_ and 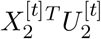 of size *n*_2_ × *k* for *X*_2_.

We set *l* as the number of pairs of canonical variables with correlation greater than a threshold λ, or 1 in the case that no pairs have this correlation. In practice, we set λ to .95 and found this threshold using cross-validation (Supplementary Methods; Section S6). By finding the sites that are best approximated by a low-rank, correlated transformation, we therefore assume that the sites with the highest scores will be representative of features that are functionally shared (i.e. correlated) between the datasets. This step is analogous to one taken by ReFACTor [24], only that we leverage the *correlated* subspace of the two datasets rather than a *variable* subspace of one dataset. Though we emphasize that *CONFINED* can be used for general sources of global biological variation, for the purpose of comparing a single use-case of *CONFINED* to other methods, we empirically fit a rule for selecting the optimal *t* for cell-type composition in whole-blood datasets as a linear function of the number of individuals in *X*_1_ and *X*_2_ (Supplementary Methods; Section S6).

*CONFINED* is available as an R package at https://github.com/cozygene/CŪNFINED. The calculations in the R package were optimized with C++ code using `Rcpp` and `RcppArmadillo`. Also included with the package is an ultra-fast function for performing CCA.

### Simulations

We evaluated the performance of *CONFINED* using a simulated study. For the simulations, we generated 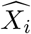 for every dataset *X_i_*:

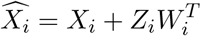

Where *Z_i_* is a random matrix of “scores” of size *m* × *r* with every entry *Z_jk_* drawn from the standard normal distribution and *W_i_* is a matrix of “weights” of size *n_i_* × *r* where every entry *W_jk_* is drawn from the standard uniform distribution and each column 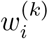 is standardized to have norm 1.

In doing so, we add some structured, normally distributed noise that is specific to each dataset. By varying the number and length of the weight vectors 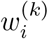, we can also control the rank and magnitude of the structured noise. Intuitively, this noise emulates technical variation, as each dataset will have its own unique set of weight vectors. For further details, see Supplementary Methods; Section S5.

### Permutation testing

To validate the enrichment results reported by `missMethyl` [47], we performed permutation testing. `missMethyl` takes as input a set (i.e. sample) of CpG sites used to test for enrichment of gene ontology pathways, along with the population from which the sample of CpG sites was chosen. For the purpose of the permutation tests, our sample of CpG sites consisted of the top *t* sites reported by *CONFINED,* and the population of CpG sites was made up of the *m* sites in the input matrices. For each number of sites *t*, we ran `missMethyl` 1000 times, using a random selection of *t* sites from the *m* sites of the input datasets at each iteration. We then compared the permutation p-values to the p-values from using the top *t CONFINED* sites. For further information, see Supplementary Methods; Section S7.

### Datasets

Throughout our experiments, we used publicly available data generated from the Illumina Infinium Human Methylation 450k chip. Our analyses focused on four whole-blood datasets and one brain-tissue dataset: (1) an analysis of Rheumatoid arthritis patients and controls with 659 individuals from Liu et al. (GSE42861) [40] (2) a study of aging with 656 individuals from Hannum et al. (GSE40279) [41] (3-4) analysis and re-analysis of schizophrenia with 847 and 675 samples from Hannon et al. (GSE80417, GSE84727) [54] and (5) a dataset from Lunnon et al. with brain tissue from 122 individuals that was used to study Alzheimer’s disease (GSE59685) [42].

The whole-blood datasets were preprocessed following guidelines suggested by Lehne et al. [55]. Using the R package `minfi` [56], we obtained and subsequently preprocessed the raw IDAT methylation files from the Liu et al. and Hannon et al. datasets. As there was no supplied IDAT file for the dataset of Hannum et al., we simply used their published intensity values. Following the guidelines of Lehne et al., we first removed single nucleotide polymorphism markers (total of 65) then applied the Illumina background correction to the obtained intensity values treating autosomal and sex chromosomes separately. We set our p-value detection threshold to 10^-16^ and set the probes whose p-values did not fall below this threshold as having missing values.

Further, we normalized the whole-blood data using quantile normalization of the intensity values, subdivided by probe type, probe sub-type, and color channel. After finalizing the intensity levels, we calculated beta-normalized methylation levels for each probe. Probes that had more than 10% of their values missing were discarded from the datasets, and the remainder of missing values were imputed using R package `impute`. Additionally, following [27], we used GLINT [57] to remove polymorphic and cross-reactive sites [58] as well as sites from non-autosomal chromosomes.

The brain dataset from Lunnon et al. was already preprocessed using the function *dasen* from R package `wateRmelon` [59]. Notably, this function also operates on the raw intensity to generate normalized beta values and uses similar preprocessing steps, including quantile normalization and the removal of single nucleotide polymorphisms. As *CONFINED* takes as input matrices with the intersection of CpG sites in two datasets, the brain dataset was also analyzed with the removal of polymorphic and cross-reactive sites as well as sites from non-autosomal chromosomes.

Additionally, we removed from our analyses outliers and samples with missing information about their sources of variability. Samples whose principal components scores were over four standard deviations away from the mean were excluded, which led to us removing six samples from the Hannum et al. dataset and two samples from the Liu et al. dataset.

We also followed filtering procedures from other works that also used the same datasets, including the removal of consistently methylated or unmethylated sites [24,27]. Prior to running any analyses, we filtered out methylation sites with standard deviation less than .02. After all preprocessing steps the dataset from (1) Liu et al. had 376021 sites and 658 individuals, (2) Hannum et al. had 382158 sites and 650 individuals, (3) Hannon et al. 381338 sites and 638 individuals, (4) Hannon et al. 382158 sites and 665 individuals, and (5) Lunnon et al. 485577 sites and 451 individuals.

## 4 Discussion

Here, we propose *CONFINED*, a sparse-CCA-based method to capture biologically replicable signal by leveraging shared structure between datasets. Specifically, we showed its use and improved accuracy over other methods in the context of capturing cell-type composition between datasets of the same tissue type. We also showed how it can be used to capture other sources of biological signal shared across datasets. Moreover, we provide evidence that *CONFINED* can be used as a feature selection mechanism, prioritizing features that are functionally shared between datasets.

Across several datasets we demonstrated that *CONFINED* accurately captured global biological sources of variability. In the case of cell-composition, the components produced by *CONFINED* better captured cell-type composition across all cell-types in methylation datasets (of the same tissue-type) than previous reference-free methods that were designed for capturing signal from cell-type composition. Additionally, *CONFINED*’s components captured other replicable sources of variability such as age and sex. While cell-type composition was better captured when using a pair of datasets of the same tissue-type, we note that other biological factors may be better captured when pairing two datasets of different tissue types. Our results provide grounds for *CONFINED* as a means to capture replicable signal from biological sources across datasets.

Additionally, *CONFINED* is robust to technical variability. Through simulations, we demonstrated that *CONFINED* accurately captures biological signal in the presence of strong, dataset-specific technical noise. Other methods that leverage decompositions of single matrices produced components corresponding to the simulated technical noise (Supplementary Methods; Section S5), but the components produced by *CONFINED* were unaffected by the simulated noise. Therefore, leveraging *multiple* datasets through *CONFINED* can provide researchers a way to robustly account for signal arising from technical variation.

Though we learned a linear rule for selecting the sparsity parameter (i.e. the number of features) in the specific case of capturing cell-type composition in methylation whole-blood datasets (Supplementary Methods Figure S6), we emphasize that the selection of the sparsity parameter in other cases may be non-trivial. Evaluating *CONFINED* on multiple datasets and sources of biological variability aside from cell-type composition, we found that the optimal sparsity parameter for cell-type composition may not be optimal for other covariates of interest. For instance, with a pair of blood datasets, sex was better captured as the number of features increased. This may be due to the fact that specific biological functions—such as the immune response—may be confined to several thousand methylation sites, whereas changes in methylation patterns due to more broad characteristics—such as age or sex—are more minute, and thus require more information or sites to capture. We therefore suggest future investigations take place and considerations about underlying biology be taken into account for selecting the optimal sparsity parameter for biological signal aside from cell-type composition.

We also showed the utility of *CONFINED* as an unbiased way of selecting informative and potentially biologically relevant methylation sites. Intuitively, as CCA finds shared structure between datasets, this structure should be reflective of biological mechanisms that are common to a pair of datasets. In our experiments, *CONFINED* found methylation sites that capture the shared variability across different blood tissues, and this set of sites was significantly enriched for immune function. Similarly, for the brain-blood pair, we observed enrichment for some immune and hematopoietic function, but the enrichment was generally not significant. Thus, our results suggest that our feature-selection method may be useful in highlighting pathways that are similar across two datasets.

A similar concept to *CONFINED* has been previously introduced in the context of singlecell RNA-sequencing by Butler et al. [53]. However, mathematically, the problem Butler et al. solve is different as the number of “individuals” (in their case, cells) in single-cell RNA is much larger than the number of features (genes), whereas in our setting, the number of individuals is much smaller than the number of features (methylation sites). Moreover, we show that a simple application of CCA does not suffice in the case of methylation, and thus *CONFINED* performs feature selection prior to performing CCA. In other words, *CONFINED* utilizes sparsity.

In summary, our results suggest that *CONFINED* will be a useful tool in capturing effects of biological variability as well as highlighting shared cellular mechanisms across multiple datasets. The components from *CONFINED* can be used in downstream analyses that wish to model only the biological signal of a methylation dataset or to include certain biological signals as confounders in statistical analyses. We suggest future research into the selection of t, the number of informative sites to use for recovering signal for specific biological factors, as well as research into which pairs of phenotypes or datasets may be useful in extracting signal for specific biological drivers of variability.

## Supplementary Methods and Information

### S1 A brief tutorial on canonical correlation analysis

In the simplest terms, CCA attempts to maximize the correlation of two matrices via linear transformations. CCA takes as input two matrices *X*_1_ of dimension *n* × *m*_1_ and *X*_2_ of dimension *n* × *m*_2_ where *n* > *m*_1_ and *m*_2_. In other words, both matrices have the same number of rows but not necessarily the same number of columns. CCA then attempts to find *m*_1_- and *m*_2_-length vectors *a*_1_ and *a*_2_, such that the correlation of *X*_1_*a*_1_ and *X*_2_*a*_2_ is maximized:

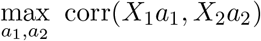

To produce *a*_1_ and *a*_2_, we first obtain vectors *b*_1_ and *b*_2_, the eigenvectors corresponding to the largest eigenvalues of the following matrices (where *X*_1_ and *X*_2_ are column-centered):

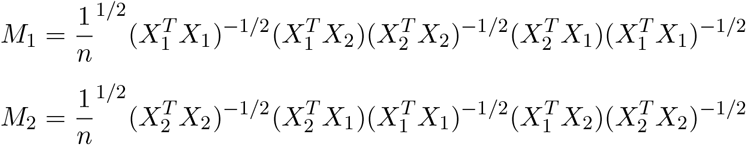

The vectors *a*_1_ and *a*_2_ are then obtained from a simple change of basis of *b*_1_ and *b*_2_ respectively:

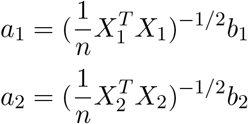

The products *X*_1_*a*_1_ and *X*_2_*a*_2_ are referred to as the first canonical variables of the input matrices, and we let *u*_1_ = *X*_1_*a*_1_ and *u*_2_ = *X*_2_*a*_2_. CCA can produce up to min{*m*_1_, *m*_2_} pairs of canonical variables from the remaining eigenvectors, however, the first pair of canonical variables (corresponding to the largest eigenvalue) has the greatest correlation.

When seeking the second and subsequent pairs of canonical variables, one additional restriction is introduced—the new canonical variables must be orthogonal to all the previous ones:

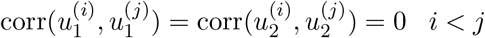

Given this constraint, the solution for the *i*^th^ pair of canonical variables conveniently follows the same formula as the first pair, only that we substitute the eigenvector corresponding to the *i*^th^ largest eigenvalue for the eigenvector corresponding to the largest eigenvalue. We then column-wise concatenate all 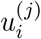 for each dataset to obtain two matrices (*U*_1_ and *U*_2_) of canonical variables of size *n* × min{*m*_1_, *m*_2_}. The canonical variables are ordered such that their correlation (which is proportional to their corresponding eigenvalue) is in decreasing order:

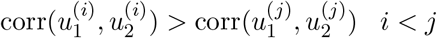

Additionally, the canonical variables have the properties that each of their variances equal 1, and the covariance of 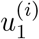 and 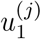 (and 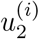 and 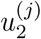) is equal to 0 when *i* ≠ *j*:

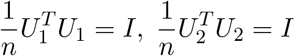

To reiterate, the basic goal of CCA is to find *a*_1_ and *a*_2_ such that corr(*X*_1_*a*_1_,*X*_2_*a*_2_) is maximized. There are min{*m*_1_, *m*_2_} such vectors for each pair of datasets, yielding min{*m*_1_, *m*_2_} pairs of canonical variables.

### S2 Feature selection

In this section, we compare the feature selection steps taken by *CONFINED* (a type of sparse CCA) and ReFACTor (a type of sparse PCA). Given input matrices of size *m* × *n*_1_ and *m* × *n*_2_, or more generally for the single-matrix decomposition case *m* × *n*.

1. Our features are selected in the following manner:

i. Obtain *U*_1_ and *U*_2_ both of size *m* × min{*n*_1_,*n*_2_} following Equations (1) and (2).
ii. Construct *Ũ*_1_ and *Ũ*_2_ from the first *l* columns of *U*_1_ and *U*_2_ respectively.
iii. Generate a low-rank approximation of each dataset:

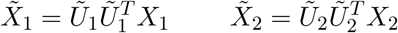
iv. For each site *j* in dataset *i* compute a score based on its correlation between itself and its low-rank approximation:

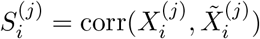
v. Rank the sites with the highest inter-dataset score:

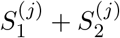
vi. Use *t* sites with the top *t* scores when performing CCA.
2. ReFACTor selects CpG sites in the following way:

i. Compute the singular value decomposition (SVD) of a matrix *X* and to obtain *V*, the left singular vectors of *X*.
ii. Construct *Ṽ* by taking the first *l* columns of *V*.
iii. Construct a low-rank approximation of *X*:

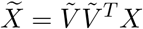
iv. Find the sites that are most correlated between the original dataset and the low-rank approximation of the dataset.
v. Use the top *t* most correlated sites for *X* when performing PCA.

Notably, the features selected by our method are important to both datasets. In both datasets, the features are well-represented in a low-dimensional, *correlated* subspace. In ReFACTor, the features are selected if they are well-approximated by the first few principal components in one specific dataset’s *variable* subspace. Performing feature selection based on a single-matrix decomposition method does not consider that some of the first few principal components in a dataset may be driven by batch effects.

### S3 Comparison of PCA and *CONFINED* using *CONFINED*-sselected features

Considering that *CONFINED* may better capture cell composition than single-matrix decompositions as it looks at characteristics shared between datasets, we provide a direct comparison of the accuracy of the top-performing single-matrix decomposition method for capturing cell-type composition, ReFACTor, and *CONFINED* using the same feature selection for both methods. We generated the rankings of features as detailed in the feature selection subsection. Even when ReFACTor uses the features generated by *CONFINED*, it captured cell-composition with lower correlation than *CONFINED* (Figure S1).

**Figure S1.**
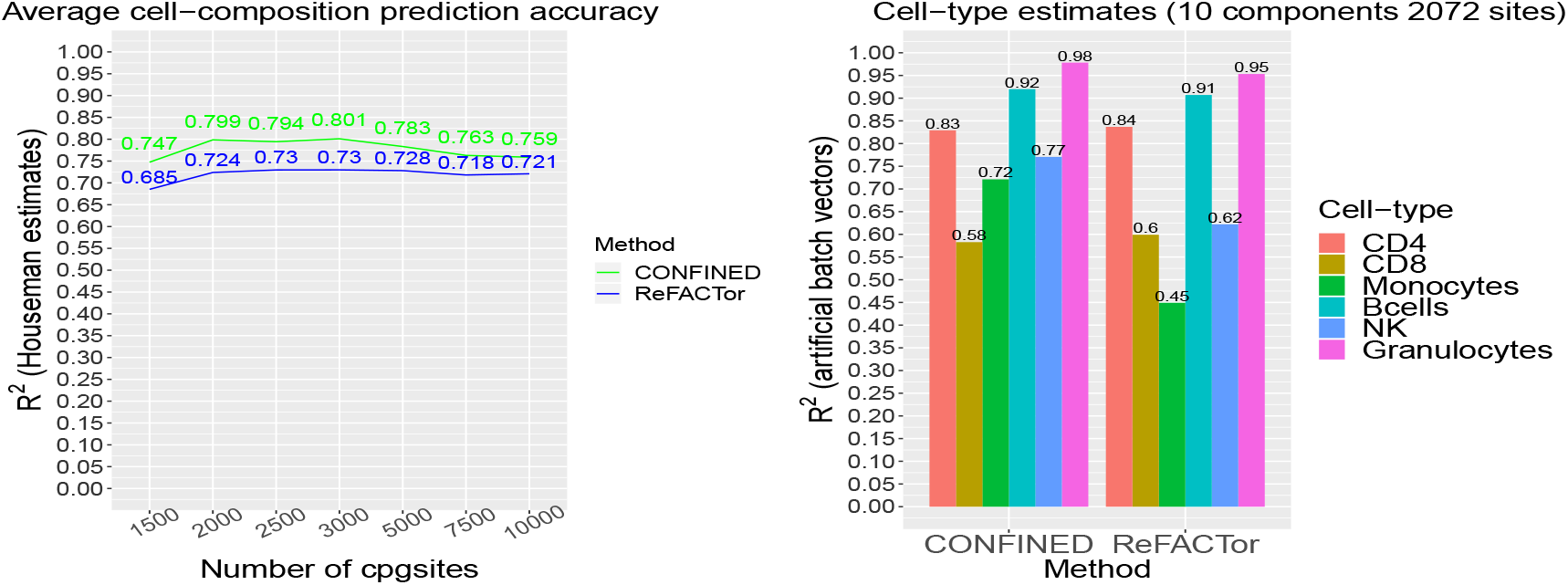
Comparison of methods based on CCA and PCA. We compared the performance of *CONFINED* and ReFACTor on dataset GSE40279 (Hannum et al. [41]) when both methods used the same features obtained in our feature selection process. On the left, the performance of each method as we varied the number of features. On the right, a comparison of the accuracy of both methods using the the sparsity parameter calculated from cross validation (Supplementary Methods Figure S6).

### S4 Single matrix decomposition on the union of two matrices

In this analysis, we evaluated the best-performing single-matrix method for capturing cell-type composition—ReFACTor—against *CONFINED*, using the union of the two input matrices to *CONFINED* as the input for the ReFACTor. To do this, we simply column-wise concatenated the individuals from one dataset to the other, so that the dimension of the new input matrix was *m* × (*n*_1_ + *n*_2_). We ran ReFACTor while also supplying it with a covariate as the `covfile` argument a vector that indicated from which dataset the individual originated (e.g. 0 for dataset_1_ and 1 for dataset_2_). We ran *CONFINED* using the learned rule for selecting the sparsity parameter since the experiment was evaluating each method’s ability to capture cell-type accuracy.

**Figure S2.**
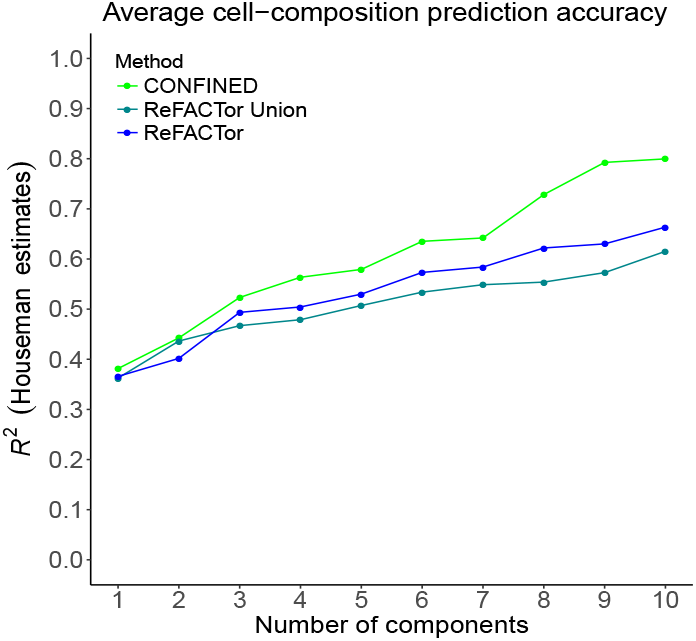
Single-matrix method using the union of two matrices. In this experiment we used datasets GSE40279 (Hannum et al. [41]) and GSE42861 (Liu et al. [40]). In green, the cell-type composition accuracy of *CONFINED* for dataset GSE40279, in turquoise the accuracy of ReFACTor when using as input the union of the two datasets, and in blue the performance of ReFACTor when just using GSE40279.

### S5 Batch effect simulations

Consider one model of principal components analysis:

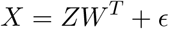

Where *X* is a data matrix of size *n* × *p, W* is an *p* × *k* matrix containing the *k* principal components of *X* (eigenvectors of the covariance matrix of *X*), *Z* is a *n* × *k* matrix of scores for each principal component, and *͞* is an *n*-length vector containing noise. Intuitively, by finding the eigenvectors corresponding to the top *k* eigenvalues of the covariance matrix of *X*, we are finding the directions that explain the most variance in the data. While we might expect that cell counts or some other phenotype might be driving the variance of methylation data, variance in biological data is often confounded by different measurement protocols or human error—in other words, batch effects [9]. Therefore, the top *k* directions of variance in a dataset may correspond to batch effects, or the observed variance in the data may simply by due to different protocols used to produce the data. For our simulations, we generated noise for each dataset *X_i_* following the previously described structure:

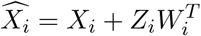

Where *Z_i_* is a random matrix of “scores” of size *m* × *r* with every entry *Z_jk_* drawn from the standard normal distribution and *W_i_* is a matrix of “weights” of size *n_i_* × *r* where every entry *w_jk_* is drawn from the standard uniform distribution and each column 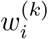 is standardized to have norm 1.

In doing so, we add some structured, normally distributed noise that is specific to each dataset. By varying the number and length of the weight vectors 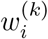, we can also control the rank and magnitude of the structured noise. Intuitively, this noise emulates technical variation, as each dataset will have its own unique set of weight vectors.

**Figure S3.**
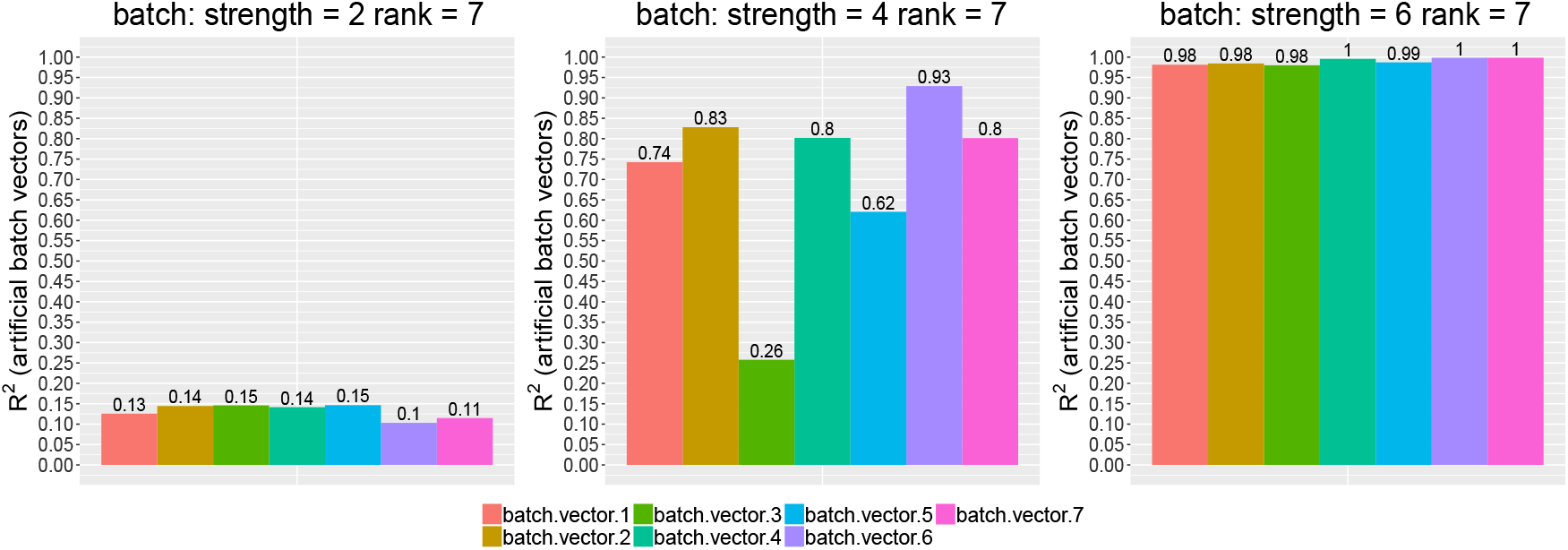
Batch-effect signal captured by ReFACTor. Here, we examine what portion of the artificial noise is captured by ReFACTor. After adding rank-7-structured noise with different strengths to each dataset, we ran ReFACTor with default settings and examined the correlation between the top 7 ReFACTor components and noise vectors.

To elucidate the consequences of the simulated batch effects on PCA-based methods, we examined the correlation of ReFACTor’s components and the simulated weight vectors. Regressing the artificial noise vectors onto the ReFACTor components, we observe high *R*^2^ values. Additionally, if the noise we introduced had rank *k* and large strength (norm), it was captured by exactly the top *k* ReFACTor components. In cases where the norm of the weight vector was relatively low, ReFACTor’s components still captured some of the signal corresponding to the batch effects. These results emphasize that single-matrix decomposition methods may produce components whose signal also includes noise from technical variation.

**Figure S4.**
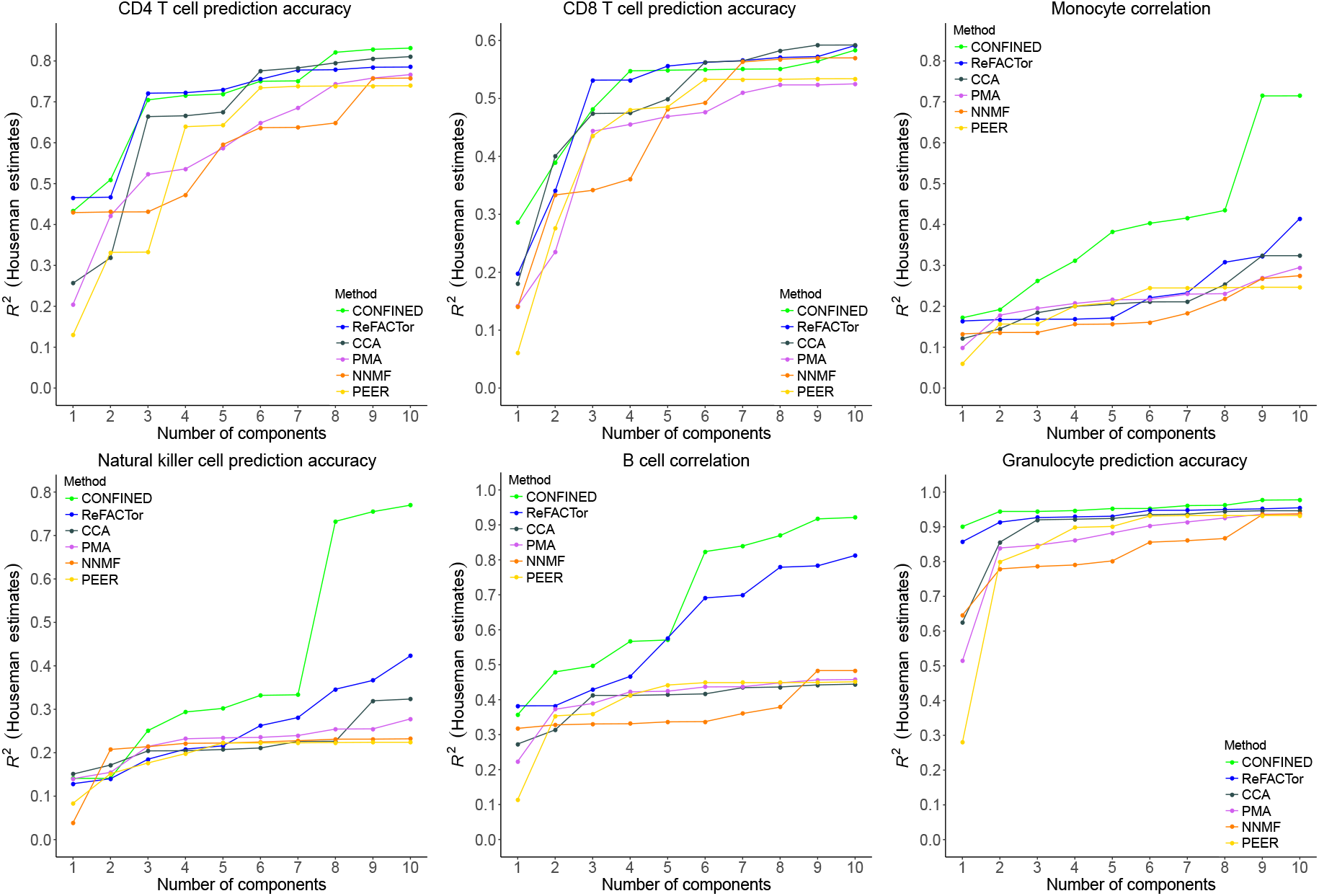
Evaluation of the ability of *CONFINED* and previous methods to capture cell-type composition. Here, we compare *CONFINED* and previous reference-free methods to capture Houseman estimates for 6 immune cell-types in a whole-blood dataset from Hannum et al. [41] using up to 10 components for each method. For *CONFINED,* we paired the whole-blood dataset from Hannum et al. with the whole-blood dataset of Liu et al. [40].

**Figure S5.**
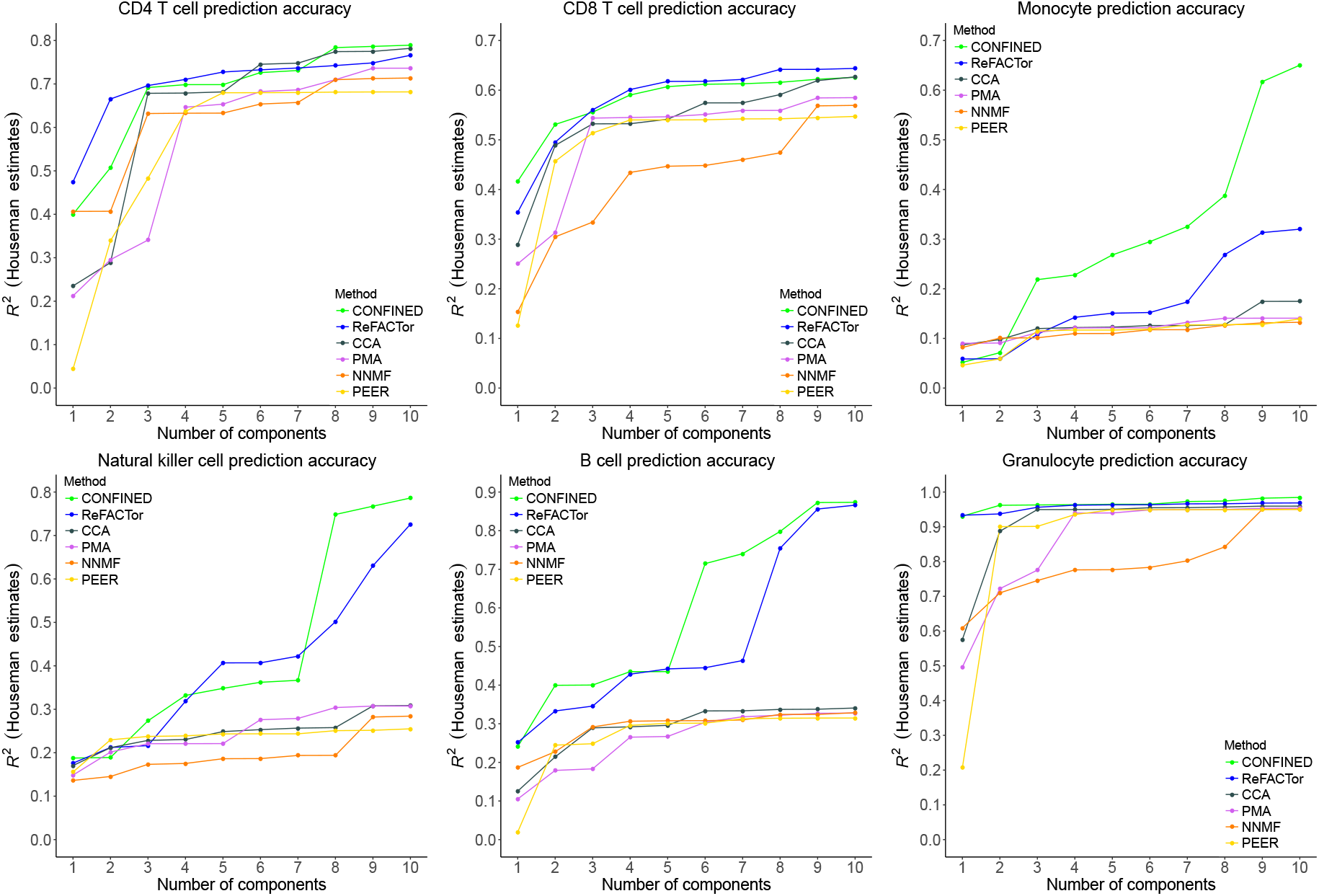
Evaluation of the ability of *CONFINED* and previous methods to capture cell-type composition. Here, we compare *CONFINED* and previous reference-free methods to capture Houseman estimates for 6 immune cell-types in a whole-blood dataset from Liu et al. [40] using up to 10 components for each method. For *CONFINED,* we paired the whole-blood dataset from Liu et al. with the whole-blood dataset of Hannum et al. [41].

### S6 Cross validation for cell-type composition

Notably, our method has two hyper-parameters, *t* the number of sites to include, and *l* the rank of the transformation used to obtain the informative sites. In this section, we explain how we chose values for both of the hyper-parameters. As a reminder, we pick the sites most correlated between their original data matrix and their low-rank approximation, e.g. *X_i_* and 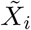 (where 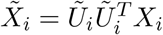). We will first explain how we choose *l*.

i. Perform CCA on the input matrices
ii. Define λ to be some threshold [0,1]
iii. Set *l* to the number of canonical variables of *U*_1_ and *U*_2_ that have correlation ≥ λ, i.e. the number of pairs of columns of *U*_1_ and *U*_2_ whose correlation is greater than or equal to λ
iv. If no canonical variables have correlation ≥ λ, set *l* = 1.

We next detail our cross-validation process. We will assume we have *l* and a ranked list of our features:

1. First, we store a partition of the data for validation purposes.

i. Hold out one third of the sites of each matrix, storing *X*_1vaiidate_ and *X*_2vaiidate_, size 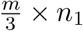 and 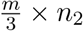 respectively
2. Using the remaining two thirds of the data, 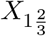 and 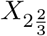, we will perform training and testing procedures.

i. Randomly split the input matrices 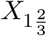 and 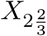 into two halves: *X*_1train_, and *X*_1test_, and *X*_2train_, and *X*_2test_ such that each matrix has 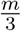 sites and their corresponding *n_i_* individuals.
ii. On the train partitions of the data, run *CONFINED* using the top *t* features to obtain *A*_1train_ and *A*_2train_, the canonical loadings for *X*_1train_ and *X*_2*train*_ respectively.
iii. Find the top *t* sites of the test data partitions and subset the test data partitions to size *t* × *n_i_* where *n_i_* is the number of individuals in that dataset.
iv. Using *A*_1train_ and *A*_2train_, obtain the *t* × *n_i_* canonical variables *U*_1test_ and *U*_2test_: *X*_1test_ *A*_1train_ and *X*_2test_ *A*_2train_ respectively.
v. Use 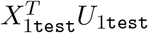 and 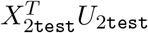 to predict cell-type composition for each individual in the test partition of the dataset.
3. After learning the optimal parameters *t*\* and *l*\*, perform our method on the validation partition of the datasets.

In this setting, we essentially learn the axes of the most correlated space for the sites of the `train` datasets, and then leverage this space on the `test` datasets to estimate cell-type composition. The canonical weights used for each test partition were obtained without using data from any of the samples in the `test` partitions.

We performed our method on `train` and `test` partitions of the data (**2** above) while varying both the value of *t* and the threshold λ. For each combination of *t* and λ, we randomly split the data 10 times, then took the average of the *R*^2^ value when using the first 10 components to capture cell-type composition as a metric of accuracy. Regressing *t* of the best performing set of hyperparameters onto the number of individuals in the datasets, we learned a rule for selecting *t* in the case of predicting cell-composition in a pair of whole-blood datasets:

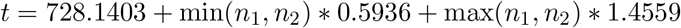

**Figure S6.**
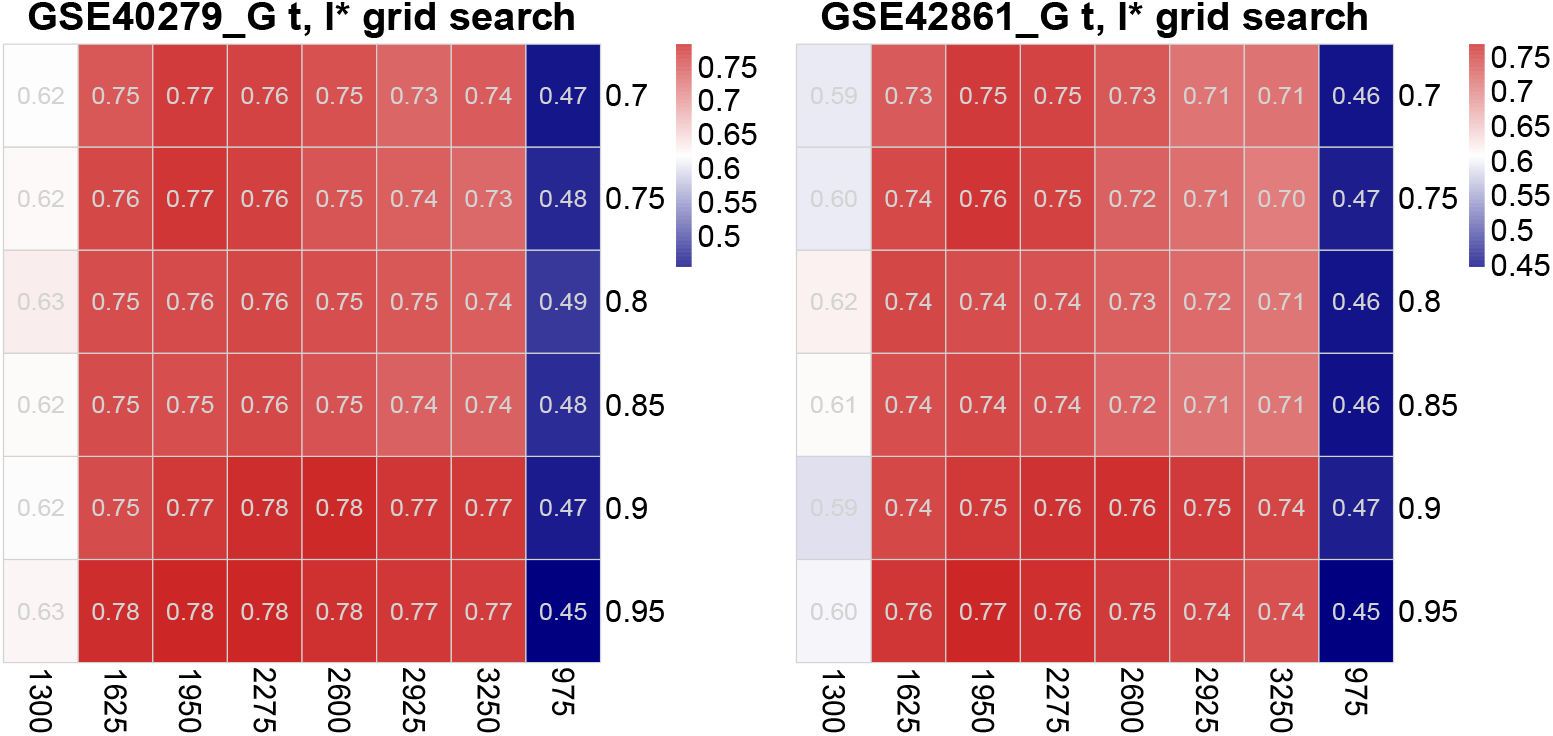
*R*^2^ of the testing partitions of the data through varying *t* and λ. We varied the number of sites to use (bottom axis) and the correlation threshold (side axis) of canonical variables to include in the feature selection step of our method. Each row is a pair of datasets used as input to our method. The number of individuals in datasets GSE40279 (Hannum et al. [41]) and GSE42861 (Liu et al. [40]) was 650 and 658 respectively.

### S7 Permutation testing

To validate the enrichment results reported by `missMethyl`, we performed permutation testing. `missMethyl` takes as input a set (i.e. sample) of CpG sites used to test for enrichment of gene ontology pathways, along with the population from which the sample of CpG sites was chosen. For the purpose of the permutation tests, our sample of CpG sites consisted of the top *t* sites reported by *CONFINED,* and the population of CpG sites was made up of the *m* sites in the input matrices. In this context, we varied t, the number of features to use, and compared the enrichment p-values when using the top *t* features sorted by *CONFINED* and *t* randomly selected features. We specifically compare the p-values of the top three most enriched pathways when using *t CONFINED* sites and the single most enriched pathway when using *t* randomly selected sites from the size *m* CpG population. The number of features we tested ranged from 1000 to 10000 with a step size of 1000, and we performed 1000 permutations at each number of features. In this experiment, we focused on a blood-blood pair of datasets (Liu et al. [40], Hannum et al. [41]).

**Figure S7.**
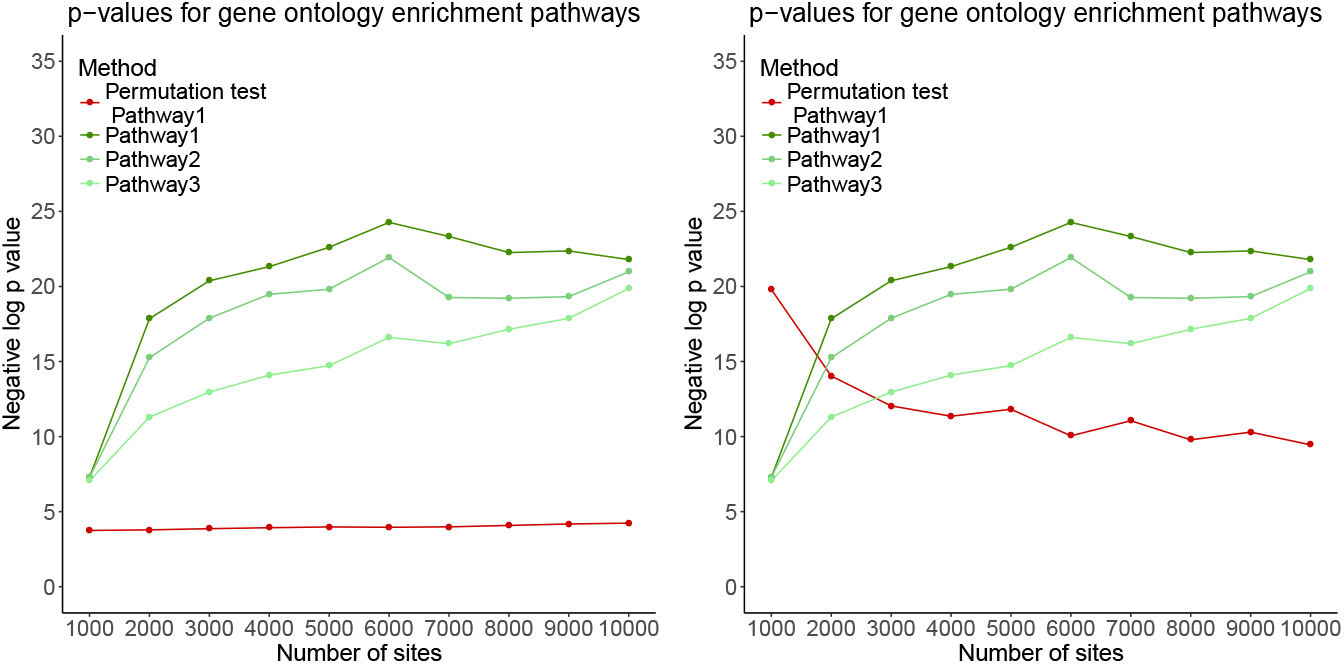
Permutation testing for gene ontology pathway summary statistics. In green are the top three pathways when choosing the top *t CONFINED*-ordered sites. The red line indicates the average (left) and minimum (right) of 1000 p-values of the top ontology pathway from *t* randomly selected sites.

### S8 Measured sources of variability across methods

In this section, we compare the performance of *CONFINED* to previous methods designed to capture cell-type composition in methylation datasets. Notably, the aim of several of the previous methods (ReFACTor, NNMF) is simply to capture cell-type composition accuracy, and they do not have an emphasis on other global sources of variation such as age, sex, or environment. For *CONFINED*, we used as input datasets from Hannum et al. [41] and Liu et al. [40]. All other single-matrix decomposition methods used just the dataset from Liu et al. as input. We used 10 components from each method to predict each source of variability in the Liu et al. [40] dataset. *CONFINED* greatly outperformed other methods when capturing both age and sex and performed best for these factors using a large number of features (i.e. low sparsity). This may be an indicator that changes in observed methylation signal due to age or sex are rather subtle, and require many sites to capture. Interestingly, when *CONFINED* used 5000 sites, age was captured with much greater accuracy than sex.

**Figure S8.**
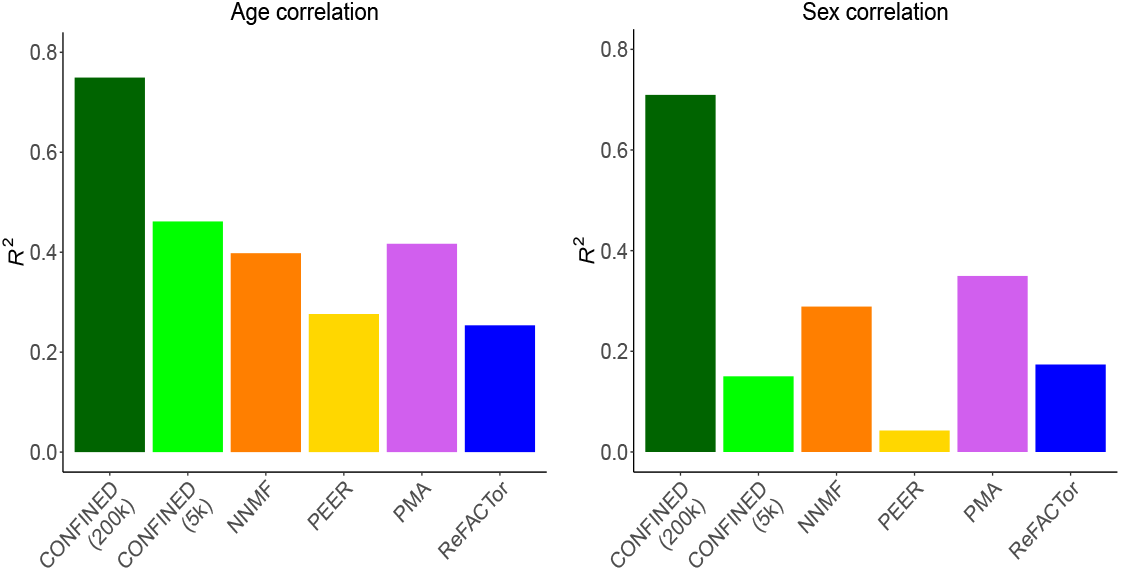
Correlation of measured sources of global variation with components from *CONFINED* and single-matrix methods. Here, we compare *CONFINED* and previous reference-free methods to capture measured sources of global variation in a whole-blood dataset from Liu et al. [40].

**Figure S9.**
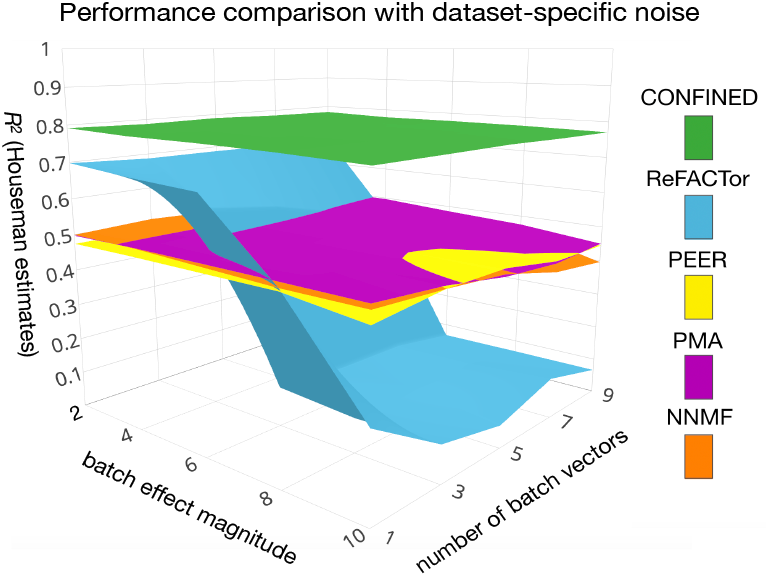
Capturing cell-composition in the presence of simulated technical noise. We added simulated batch effects to the whole-blood datasets of Liu et al. [40] and Hannum et al. [41] and compared the ability of *CONFINED,* ReFACTor [24], PEER [38], PMA [36], and NNMF to capture cell-type composition in whole-blood. Here, we show the results of the Liu et al. dataset.

